# Fast recovery of disrupted tip links induced by mechanical displacement of hair bundles

**DOI:** 10.1101/2020.10.02.324111

**Authors:** R. G. Alonso, M. Tobin, P. Martin, A. J. Hudspeth

**Author notes:** Corresponding author: Dr. A. J. Hudspeth, Laboratory of Sensory Neuroscience, The Rockefeller University, New York, NY 10065. Author contributions: R.G.A, M.T., P.M., and A.J.H. designed the research; R.G.A and M.T. performed the research and analyzed the data; R.G.A, P.M., and A.J.H. wrote the paper. Perelman School of Medicine, University of Pennsylvania, Philadelphia, PA, USA.

## Abstract

Hearing and balance rely on the capacity of mechanically sensitive hair bundles to transduce vibrations into electrical signals that are forwarded to the brain. Hair bundles possess tip links that interconnect the mechanosensitive stereocilia and convey force to the transduction channels. A dimer of dimers, each of these links comprises two molecules of protocadherin 15 (PCDH15) joined to two of cadherin 23 (CDH23). The “handshake” that conjoins the four molecules can be disrupted *in vivo* by intense stimulation and *in vitro* by exposure to Ca^2+^ chelators. Using hair bundles from the rat’s cochlea and the bullfrog’s sacculus, we observed that extensive recovery of mechanoelectrical transduction, hair-bundle stiffness, and spontaneous bundle oscillation can occur within seconds after Ca^2+^ chelation, especially if hair bundles are deflected towards their short edges. Investigating the phenomenon in a two-compartment ionic environment that mimics natural conditions, we combined iontophoretic application of a Ca^2+^ chelator to selectively disrupt the tip links of individual frog hair bundles with displacement clamping to control hair-bundle motion and measure forces. Our observations suggest that, after the normal Ca^2+^ concentration has been restored, mechanical stimulation facilitates the reconstitution of functional tip links.

**Significance Statement:** Each of the sensory receptors responsible for hearing or balance—a hair cell—has a mechanosensitive hair bundle. Mechanical stimuli pull upon molecular filaments—the tip links—that open ionic channels in the hair bundle. Loud sounds can damage hearing by breaking the tip links; recovery by replacement of the constituent proteins then requires several hours. We disrupted the tip links *in vitro* by removing the calcium ions that stabilize them, then monitored the electrical response or stiffness of hair bundles to determine whether the links could recover. We found that tip links recovered within seconds if their ends were brought back into contact. This form of repair might occur in normal ears to restore sensitivity after damage.

---

Our ability to hear sounds and detect accelerations stems from the capacity of specialized cells to transduce mechanical energy into electrical signals that are interpreted by the nervous system. Transduction is accomplished by mechanoreceptors known as hair cells that—with subtle morphological and physiological variations—operate in fundamentally the same way among all vertebrates (1)(2)(3).

Each hair cell detects mechanical stimuli with an organelle, the hair bundle, composed of tens to hundreds of rod-like, actin-filled stereocilia that protrude from the cuticular plate, an actin-rich structure at the cell’s apical surface (4). These stereocilia are arranged in a staircase, increasing in length monotonically towards a single true cilium termed the kinocilium (5). In the mammalian cochlea, the kinocilium degenerates after the hair bundle has developed, but the organelle persists in hair cells from the mammalian vestibular system and those from other vertebrates. Upon deflection of the hair bundle, the stereocilia pivot about their insertions into the cuticular plate; the resultant shear between contiguous stereocilia modulates the extension of elastic elements that are coupled to mechanoelectrical-transduction channels, eliciting an electrical response (6, 7). By controlling the resting tension of these gating springs, adaptation motors set the channels’ open probability and thus regulate the sensitivity of mechanoelectrical transduction (8)(9)(10).

Each stereocilium bears an oblique filament—the tip link—that connects its tip to the flank of a neighbor in the taller stereociliary row and is thought to be a component of the gating spring (11, 12)(13). A tip link consists of a parallel homodimer of cadherin 23 (CDH23) in its upper two-thirds and a parallel homodimer of protocadherin 15 (PCDH15) in its lower third (14). Each of the 38 unique extracellular cadherin domains in the two proteins is stabilized in part by binding of Ca^2+^ to sites at one or both of the domain’s ends (15).

Ca^2+^ also stabilizes the molecular handshake that interconnects the proteins at their amino termini (16). Exposure of a hair cell to a Ca^2+^ chelator for as little as a few seconds disrupts these interactions and terminates mechanoelectrical transduction (17, 18). Although the handshake interaction might potentially be regenerated after the restoration of Ca^2+^, there are at least three reasons why such recovery might not occur. First, each tip link in a resting hair bundle bears a resting tension on the order of 10 pN (8)(19). When a link is severed, the two ends are expected to undergo elastic retraction from one another. The second issue is that the tension in intact tip links pulls a hair bundle toward its short edge by exerting force against the flexion of the stereociliary pivots. When all the tip links of a frog’s hair bundle are disrupted, the bundle can lunge more than one hundred nanometers in the positive direction (17). The stiffer hair bundles of the rat’s cochlea move somewhat less (8). As a result of the geometric relationship between the motion of a hair bundle and the shear between contiguous stereocilia, the larger movement would displace the separated ends of a tip link by up to 20 nm. A final possibility is that dissociated cadherins are internalized and therefore no longer available to reconstitute tip links (20). In the present investigation, we inquired whether tip links might recover if their separated ends were re-apposed, for example by deflecting a hair bundle well in the negative direction before internalization could occur.

## Results

### Rapid recovery of mechanoelectrical transduction upon hair-bundle deflection

To determine whether tip-link integrity can be restored on a short time scale after disruption, we first measured the mechanoelectrical transduction currents from outer hair cells of an excised preparation of the neonatal rat’s cochlea before and after disrupting the tip links. We used a calibrated fluid jet to deflect each hair bundle, voltage clamping to measure the transduction current, and iontophoresis to deliver ethylenediaminetetraacetic acid (EDTA) onto the hair bundle. We found that in response to a 60 Hz sinusoidal stimulus that deflected the hair bundle by approximately 100 nm, the transduction current could recover in part within 1 s after the iontophoretic pulse. In one example (Fig. 1*A*), comparison of the responses measured before tip-link disruption and after recovery indicated that the transduction current achieved 57 % of its original level. For this cell the mean and variance of the transduction current reached 72 % and 49 % of their respective control values. The time course of the recovery was roughly exponential with a time constant of 100 ms (Fig. 1*B*). Nine of the 16 outer hair cells that we examined displayed recovery of their transduction currents during sinusoidal stimulation to at least 100 pA, or 20-60 % of the original level

**Fig. 1.**
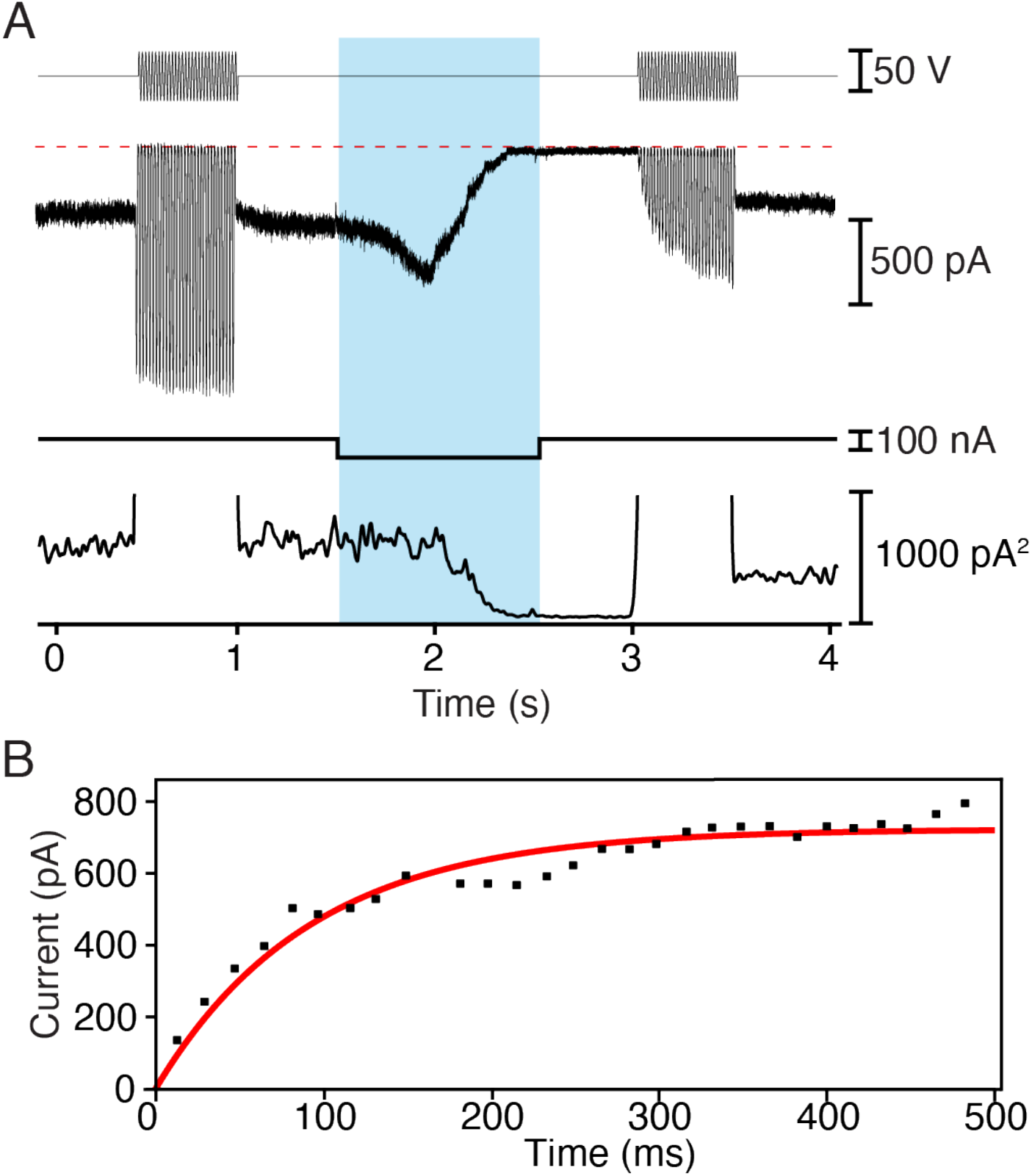
Rapid recovery of mechanoelectrical transduction in outer hair cells from the rat’s cochlea. (*A*) A fluid-jet stimulator was driven at 60 Hz with two sinusoidal stimulus trains (top trace), one before and one after an iontophoretic current pulse (bottom trace) that released EDTA. The pale blue band here and in subsequent figures delineates the period of iontophoresis. The transduction current (second trace) was initially large, but fell to nearly zero after iontophoresis before recovering more than half of its original magnitude during the second stimulus train. The variance of the transduction current (third trace) fell during iontophoresis as transduction was interrupted, but recovered partially after a second epoch of stimulation. The abscissa represents zero variance. (*B*) For the record shown in panel *A*, the recovery of the transduction current after the iontophoretic pulse followed an exponential relation (red line) with a time constant of 100 ms.

These observations could not be explained by incomplete disruption of the tip links: before the second stimulus train, both the mean and the variance of the resting current were negligible, reflecting the absence of mechanotransduction. Furthermore, the time evolution of the current during the iontophoretic pulse reflected the known response of the transduction machinery to a Ca^2+^ chelator (18)(8): the current first increased owing to a rise in tip-link tension, then fell to zero as the links were disrupted and the transduction channels closed. The variability in recovery likely reflected differences in the number of tip links that were reconstituted before the component cadherins moved away or were subducted from the membrane.

### Rapid recovery of mechanical properties upon hair-bundle deflection

Electrical recording in a low-Ca^2+^ environment that emulates endolymph is difficult. Moreover, we ascertained that iontophoresis of a Ca^2+^ chelator does not disrupt tip links in solutions with Ca^2+^ concentrations in the millimolar range. The mechanical properties of the hair bundle offered an alternative manifestation of the disruption and recovery of tip links. We therefore designed a series of protocols to examine how the process occurs. The fragility of the mammalian cochlea largely precludes an *ex vivo* preparation that recreates the ionic milieu in which the cochlea normally operates. The two-compartment preparation of the bullfrog’s sacculus, however, reconstitutes the ionic environment of hair cells and retains most of the active characteristics of the cells (21, 22). In addition, the high cohesiveness of bullfrog hair bundles ensures that they move as a unit in response to mechanical stimulation, which facilitates the interpretation of stiffness measurements (6). We therefore elected to investigate the recovery process further with that preparation.

We attached the tip of a flexible glass fiber to the kinociliary bulb and moved the fiber’s base sinusoidally at 10 Hz with an amplitude of 100 nm. We then evaluated the displacement of the hair bundle as an estimate of tip-link integrity. Tip links contribute considerably to the total stiffness of a hair bundle (8, 17–19). For hair bundles of the bullfrog’s sacculus, as an example, tip-link disruption by Ca^2+^ chelators decreases the stiffness from *circa* 1200 μN·m^−1^ to 200 μN·m^−1^ (18). The residual stiffness is attributed to the actin filaments at the stereociliary pivots, the basal insertions of the stereocilia (23).

The mechanical responses of bullfrog hair bundles were consistent with the observations from the rat’s outer hair cells. Before an iontophoretic pulse, stimulation typically resulted in sinusoidal oscillations 50 nm in amplitude. Immediately after the iontophoretic pulse, the amplitude of oscillation increased to approximately 90 nm, then progressively declined toward the initial value before reaching a plateau near 60 nm (Fig. 2*A*). The decrease in the amplitude of the oscillations followed an exponential trajectory with a time constant of about 1900 ms (Fig. 2*B*). The change in the amplitude of the oscillations implied that the stiffness of the hair bundle fell to 55 % of its control level immediately after iontophoresis. By the end of the recording, however, the bundle regained 83 % of its original stiffness.

**Fig. 2.**
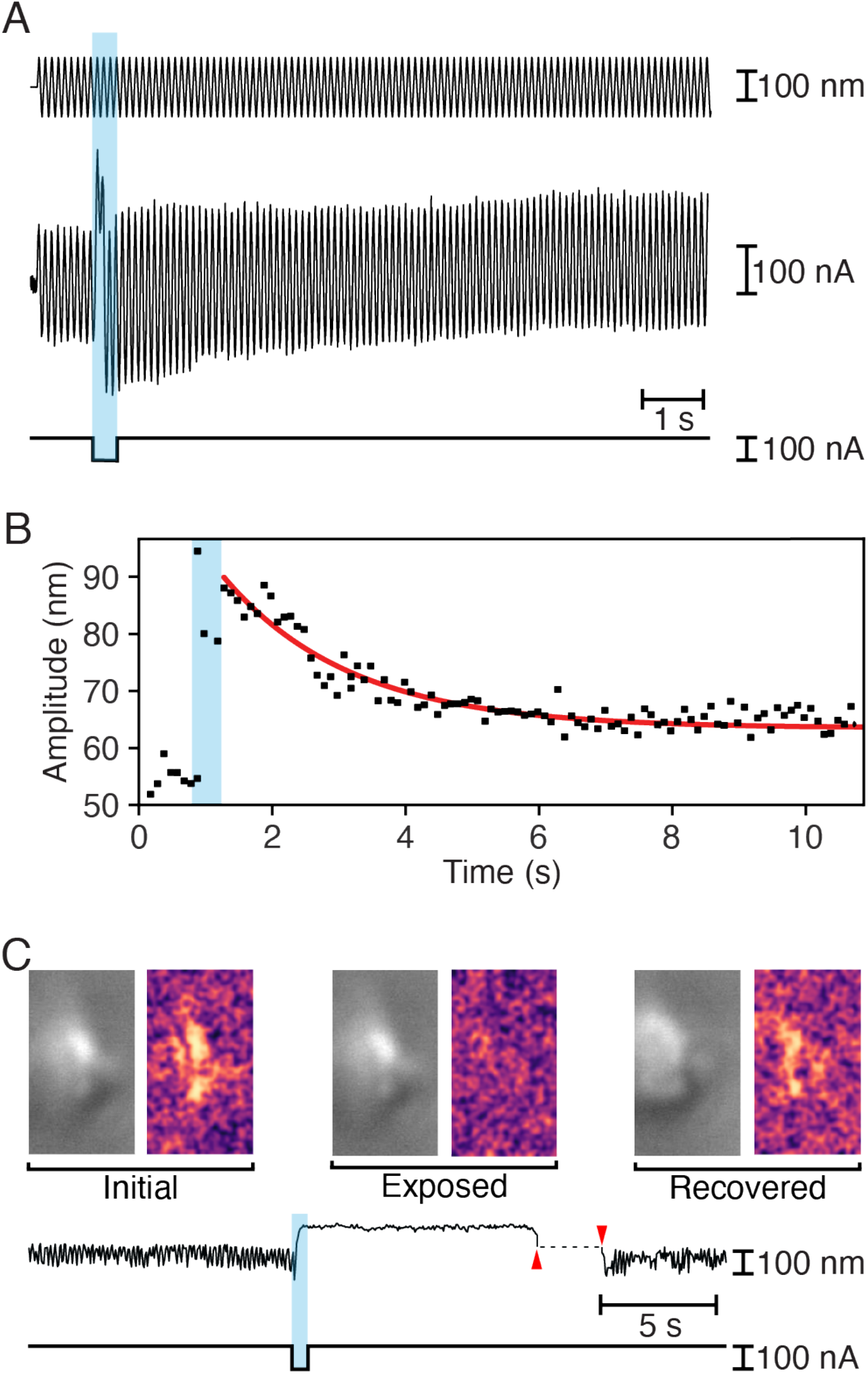
Rapid recovery of mechanical properties in hair bundles of the bullfrog’s sacculus. (*A*) While a 10 Hz sinusoidal stimulus of amplitude 100 nm (top trace) was delivered to the base of a flexible glass fiber, a pulse of iontophoretic current (bottom trace) released EDTA. The hair bundle’s movement (middle trace) increased immediately after iontophoresis, but returned toward the initial value over a few seconds. (*B*) For the record shown in panel *A*, the decline in the amplitude of hair-bundle oscillation after the iontophoretic pulse followed an exponential relation (red line) with a time constant of 1910 ms. (*C*) To the left in each pair of images in the top row are individual frames of a video of a spontaneously oscillating hair bundle (*SI Appendix*, Video S1) representing the unperturbed bundle (*Initial*), the same bundle after exposure to EDTA (*Exposed*), and finally the bundle after transient displacement in the negative direction (*Recovered*). To the right are three images, each obtained by subtracting the original frame from the subsequent frame. The *Initial* and *Recovered* images reveal spontaneous hair-bundle motion, which is absent in the *Exposed* image. The time course of the hair bundle’s position in the video (upper trace) shows suppression of the spontaneous oscillations during iontophoretic application of EDTA (lower trace) and their recovery after the bundle was pushed in the negative direction (between the red arrowheads).

When exposed to physiologically appropriate ionic solutions, hair bundles of the frog’s sacculus oscillate spontaneously (21). These movements emerge from the interplay between negative stiffness and the adaptation machinery (24). Because spontaneous oscillations require the normal operation of the transduction apparatus, disrupting the tip links would be expected to arrest the movements, which might resume if the tip links were healed.

To examine this possibility, we selected hair bundles that displayed large spontaneous oscillations that were readily recognized upon microscopic observation. Upon iontophoresis of EDTA, each hair bundle underwent an abrupt positive displacement and the oscillations stopped. Left to itself in a control experiment, the bundle remained quiescent for minutes. If instead we used a stiff glass fiber to displace the hair bundle in the negative direction for a few seconds, the oscillations promptly resumed (Fig. 2*C*; *SI Appendix*, Video S1). This result is consistent with the reconstitution of tip links and the recovery of mechanoelectrical transduction.

### Stiffness recovery as an indication of tip-link reattachment

We next used displacementclamp feedback to control the bundle’s position and to monitor the force necessary to maintain that position. With a flexible fiber attached to the kinociliary bulb of an individual hair bundle, we iontophoretically delivered EDTA to selectively disrupt the tip links. This approach allowed us to apply various displacement protocols and to assess the stiffness of a hair bundle throughout an experiment.

As expected, the displacement clamp kept the amplitude of hair-bundle movement relatively constant during sinusoidal stimulation, but the force required to do so decreased after exposure of the bundle to Ca^2+^ chelator (Fig. 3*A,B*). Because clamping was incomplete, the bundle’s excursion also increased somewhat. Both changes implied a greater compliance of the bundle owing to the disruption of tip links. For seven hair bundles from six preparations, the stiffness after chelation fell to 230 ± 16 μN·m^−1^ (mean ± SEM; Table 1), a value comparable to the stiffness of the stereociliary pivots (18). This result implies that tip-link disruption was nearly complete.

**Fig. 3.**
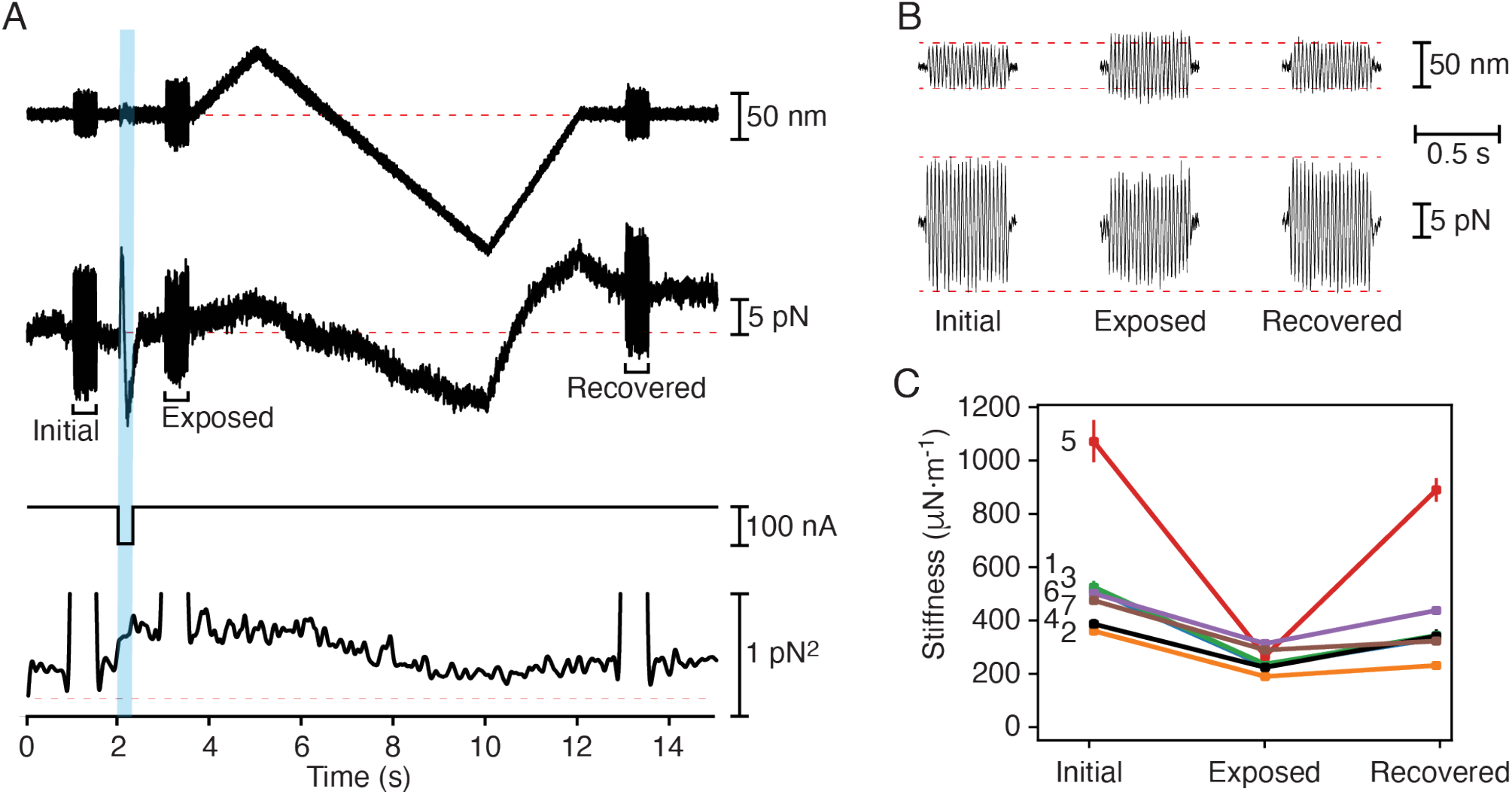
Facilitation of hair-bundle recovery in the bullfrog’s sacculus by mechanical displacement. (*A*) In a displacement-clamp experiment, a feedback system imposed a ramp displacement on a hair bundle (first trace), moving the bundle first in the positive direction to prevent prompt recovery, and then more extensively in the negative direction. At three times during this paradigm, a 500 ms epoch of ±25 nm, 40 Hz sinusoidal stimulation was superimposed on the displacement-command signal. An iontophoretic pulse (third trace) released EDTA to break tip links. The force (second trace) necessary to clamp the bundle at the outset (*Initial*) diminished after exposure to iontophoretically applied EDTA (*Exposure*) but recovered almost completely by the experiment’s end (*Recovered*). The variance of the force (fourth trace) confirmed the bundle’s softening after iontophoresis and its recovery during the negative phase of the ramp. The dashed line represents the background noise. (*B*) Enlarged records of the hair-bundle displacement (top traces) and clamp force (bottom traces) from panel *A* demonstrate that maintaining an oscillation of similar—or even greater—magnitude required less force shortly after iontophoresis. (*C*) Data from seven hair cells, which are numbered as in Table 1, reveal a significant decrease (*P* < 0.01 by a single-sided paired *t*-test) in hair-bundle stiffness after iontophoretic pulses. The stiffness then recovered significantly (*P* < 0.05 by the same test) following negative hair-bundle displacements. The bundle whose responses are depicted in panels *A* and *B* is number 4; standard deviations are shown when they exceed the size of the data points.

**Table 1.**
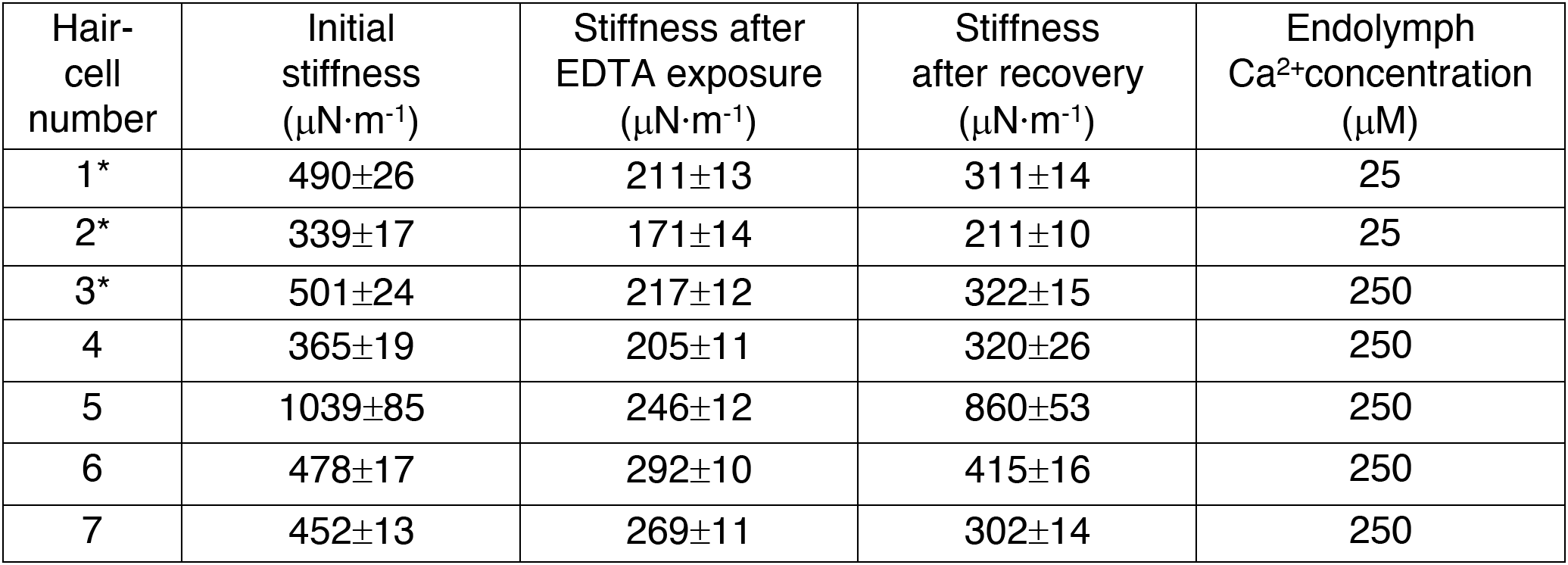
Stiffness recovery by bullfrog hair cells. Each stiffness was estimated by measuring the flexion of a flexible glass fiber attached at the hair bundle’s tip during sinusoidal stimulation. Asterisks indicate bundles subjected to step displacements; the remainder were displaced with ramps. Values are reported as means ± SDs for 21 determinations. For the entire sample, the stiffness decreased significantly after exposure with respect to the initial value (*P* < 0.007). After the negative displacements, the stiffness recovered significantly with respect to the exposed value (*P* < 0.04). If we disregard the cell (number 5) with an exceptionally high stiffness, the corresponding values show still greater significance (respectively *P* = 0.0001 and *P* < 0.002).

To foster the possible reformation of tip links, we then displaced each hair bundle as much as −150 nm with a slow displacement ramp. Moving the bundle toward its short edge—a negative stimulus—would be expected to bring the tips of successive ranks of stereocilia closer together and might therefore promote the reassociation of tip-link cadherins. Indeed, upon stimulation after the displacement, the force required during sinusoidal stimulation attained nearly its control level (Fig. 3*A*,*B*). A similar result was obtained with a step displacement (*SI Appendix*, Fig. S1 and Fig. S2). The hair-bundle stiffness therefore recovered substantially (Table 1). Comparison of the stiffnesses before and after treatment showed an average recovery of 73.5 ± 4.4 % (mean ± SEM, *N* = 7). The recovery of 81.0 ± 4.9 % (mean ± SEM, *N* = 4) during a ramp exceeded that of 63.4 ± 0.6 % (mean ± SEM, *N* = 3) for a step.

The variance in the force applied to the hair bundle provided an additional, qualitative indication of the change in a bundle’s stiffness during and after the disruption of tip links and indicated when tip-link recovery occurred (Fig. 3*A*). For the same seven hair bundles, the variance in hair-bundle force increased from a control value of 0.47 ± 0.01 pN^2^ to 0.92 ± 0.10 pN^2^ (means ± SEMs) immediately after an EDTA pulse. During the ramp protocol the variance remained high until the hair bundle was pushed in the negative direction, whereupon it fell progressively to a plateau of 0.53 ± 0.01 pN^2^ (mean ± SEM), a value slightly exceeding that prior to the iontophoretic pulse.

As a result of separation between the ends of the cadherin molecules in a disrupted tip link, recovery should be less likely if a hair bundle remains in its resting position or is moved in the positive direction. Consistent with that expectation, recovery was never observed in a hair bundle displaced in the positive direction, nor was any change noted during control experiments in which no chelator was applied (*SI Appendix*, Fig. S3). These observations confirmed that the changes in stiffness resulted from the joint action of disrupting the tip links with a Ca^2+^ chelator and displacing the stereocilia in the negative direction.

### Negative hair-bundle movement during exposure to Ca^2+^ chelators

The sequence of hairbundle forces associated with the disruption and regeneration of tip links revealed unexpected complexity in recordings from bullfrog hair cells. In six of the seven cells examined, there was a sustained positive offset of 20.1 ± 7.0 pN (mean ± SEM) at the end of the stimulation protocol with respect to the value before EDTA exposure (*SI Appendix*, Note S1). In the absence of displacement clamping, bundles displayed a corresponding offset in displacement that occured even after the kinocilium had been dissected free of the clustered stereocilia (*SI Appendix*, Note S2 and Figs. S4 and S5). The basis of this phenomenon is uncertain.

### Repeated recovery of stiffness by a hair bundle

In two experiments we examined the possibility that tip links can recover after successive treatments with Ca^2+^ chelator. By applying to the same hair bundle sequential ramp protocols separated by 10-20 s, we were able to obtain some degree of tip-link recovery after as many as three cycles of iontophoresis (Fig. 4). In each instance the recovery was partial, so the stiffness declined progressively towards the value associated with the stereociliary pivots. After the bundle’s stiffness had reached that level, no further recovery occurred. Repeated recovery of the transduction current was also observed in one hair cell of the rat’s cochlea through five cycles of iontophoresis.

**Fig. 4.**
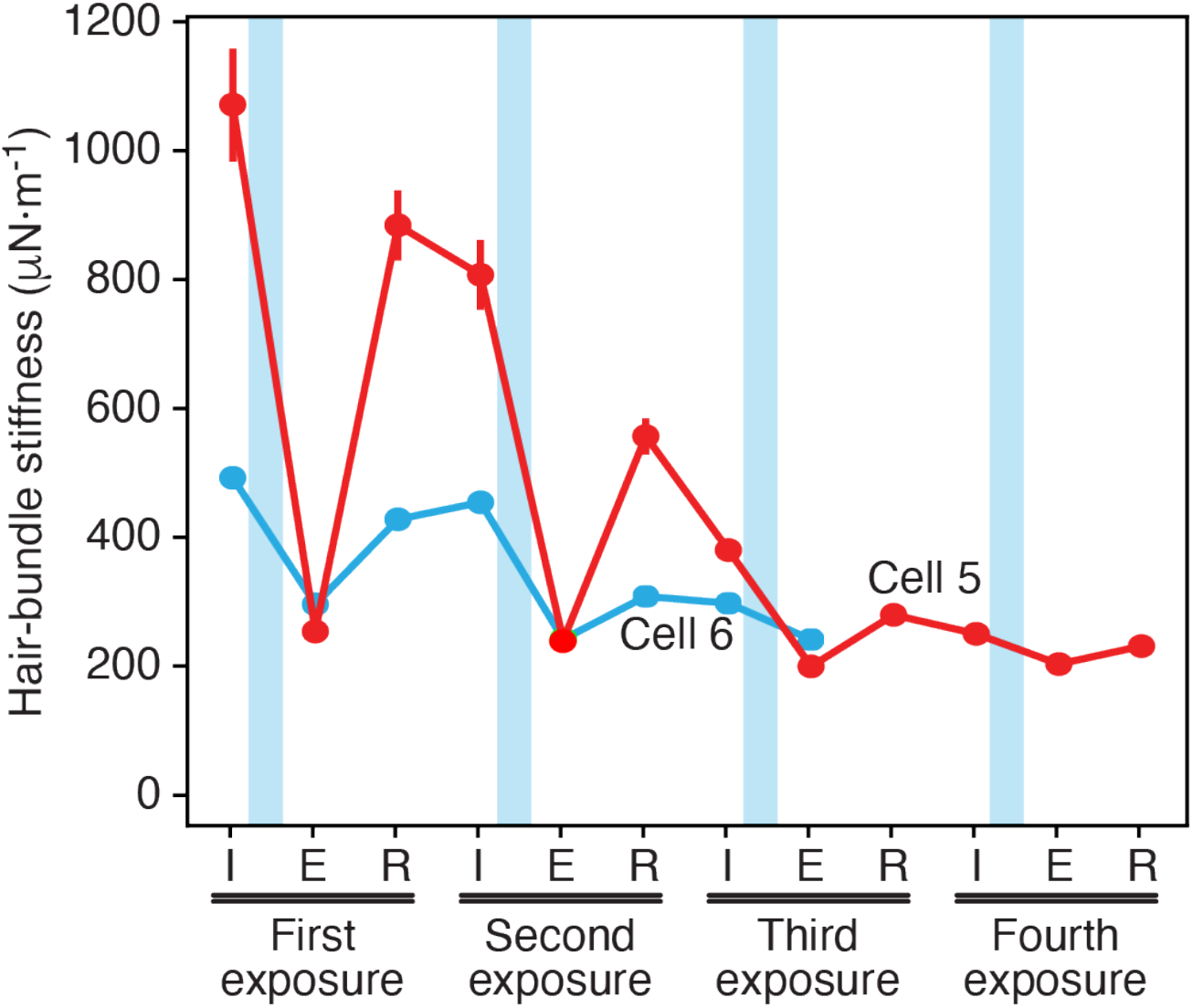
Sequential disruption and recovery of tip links in the bullfrog’s sacculus. For two hair bundles also described in Table 1, successive applications of EDTA reduced the stiffness to approximately that associated with the stereociliary pivots. When subjected to ramp displacements, one bundle recovered part of its stiffness at least three times and the other twice. The data points represent the initial stiffness (*I*), that just after EDTA exposure (*E*), and that following recovery (*R*).

### Resting tension in tip links

The resting tension of tip links was first estimated for the bullfrog’s saccular hair cells maintained in a homogeneous ionic environment (19). The present experiments afforded an opportunity to make corresponding measurements on unrestrained, oscillating hair bundles in a two-compartment chamber with more physiologically appropriate saline solutions bathing the apical and basal cellular surfaces.

In a resting hair bundle, the force exerted by tip-link tension is equal and opposite that owing to the flexion of the stereociliary pivots. By measuring the movement *X*_SP_ of each bundle upon disruption of its tip links, one may therefore calculate the average tension *t*_TL_ along the oblique axis of each tip link as

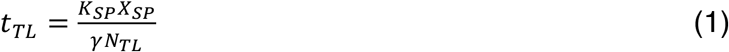

in which *K*_SP_ = 250 μN·m^−1^ represents the stiffness of the stereociliary pivots, *γ* = 0.14 the geometrical gain factor, and *N*_TL_ = 40 the number of tip links. Because a healthy hair bundle in a two-compartment chamber usually oscillates spontaneously, we measured *X*_SP_ as the bundle’s displacement from the midpoint between the positive and negative extremes of its oscillation (*SI Appendix*, Fig. S6). For 13 oscillatory hair bundles, we found *X*_SP_ = 462 ± 93 nm (mean ± SEM) and estimated the average tension in the tip links of an unstimulated hair bundle as *t*_TL_ = 20.6 ± 4.1 pN (mean ± SEM). Even in a two-compartment environment, some hair bundles did not oscillate, possibly as a result of mechanical damage or Ca^2+^ loading during the dissection in standard saline solution. Quiescent hair bundles showed smaller movements upon EDTA iontophoresis, *X*_SP_ = 187 ± 24 nm (mean ± SEM; n = 31). If the number of tip links remains constant, this implies a lower tip-link tension, *t*_TL_ = 8.3 ± 1.1 pN. Tip links in oscillatory hair bundles thus appeared to bear more than twice the tension borne by those in quiescent bundles.

## Discussion

Tip-link disruption is a well-characterized form of hair-cell damage (25). Intense hair-bundle stimulation caused by prolonged exposure to loud sounds can damage tip links *in vivo* (26), whereas Ca^2+^ chelators can dissociate them *in vitro* (17, 18). This damage can be partially reverted in 12-48 hours as the tip links regenerate and restore mechanosensitivity through a two-step mechanism (20, 27–30). Interacting PCDH15 molecules first form temporary tip links that partially restore transduction but not adaptation. The upper portions of the tip links are then replaced with CDH23 to restore normal transduction (20).

The complete replacement of tip links is both metabolically costly and relatively slow. In this study we have shown that tip links can recover within seconds after disruption by Ca^2+^ chelation. This phenomenon constitutes an unusual form of repair for a molecular lesion: disrupted links evidently reconstitute themselves from their components. One possible utility of the recovery process is the restoration of hair-bundle function after exposure to injurious events such as loud sounds (25)(26). Tip links might accordingly act as security releases that prevent more extensive damage to a hair bundle owing to overstimulation (31).

The rapidity of the recovery suggests that the phenomenon involves the reconstitution of the original tip links rather than the mobilization of stored cadherin molecules or the synthesis of new ones. There are two plausible mechanisms of recovery that are not mutually exclusive. It seems most likely that chelation disrupts the molecular handshake between PCDH15 and CDH23 dimers, and that after the restoration of Ca^2+^ the amino termini simply diffuse until they collide and reconstitute the handshake. By approximating the free ends of the PCDH15 and CDH23 dimers, pushing a hair bundle in the negative direction facilitates this process. An alternative possibility is that the handshake is never disrupted, but that the softening of tip links during Ca^2+^ chelation reflects the unfolding of extracellular cadherin domains. Such an event would profoundly affect a tip link: each 4 nm domain includes a total of about 35 nm of polypeptide chain, so the unfolding of only a few domains would result in a greatly elongated structure with a far lower stiffness. According to this model, the recovery upon the restoration of Ca^2+^ would reflect the refolding of the cadherin domains, which for PCDH15 monomers occurs within seconds in the absence of tip-link tension (13).

Under conditions in which the hair bundles oscillated spontaneously, our measured value for the positive movement of bullfrog hair bundles following EDTA iontophoresis was as much as threefold that obtained earlier (17). The difference likely reflects the fact that the present data were obtained in a two-compartment recording chamber, so that the stereocilia were bathed in low-Ca^2+^ endolymph and the cell somata in perilymph, rather than both surfaces in a homogenous medium. Calcium ions allow the adaptation motors of hair cells to slip down the stereocilia (32) and might have reduced tip-link tension in the previous study. As a consequence of the greater movements in the present experiments on oscillatory hair bundles, our estimate of the tension in individual tip links is more than twice that reported for recordings for quiescent hair bundles in a homogeneous solution (19) and resembles that for outer hair cells at the apex of the rat’s cochlea (8). Moreover, the reported value for tip-link tension is a minimal estimate: we assumed a total of 40 tip links in each hair bundle, nearly the maximum possible number for a large bundle, when in reality some bundles were smaller and some links were likely broken during dissection.

## Methods

Detailed methods are provided in *SI Materials and Methods*.

### Electrical recording from mammalian hair cells

The sensory epithelial of young rats and mature bullfrogs were maintained under microscopic observation in species-appropriate physiological saline solutions. While mechanoelectrical transduction of each outer hair cell from the rat’s cochlea was monitored by whole-cell, tight-seal recording, the hair bundle was displaced sinusoidally by fluid-jet stimulation. After tip links had been disrupted by the iontophoretic application of EDTA, the electrical response was recorded for indications of recovery.

### Mechanical recording from anuran hair cells

Each hair bundle from the frog was displaced with an elastic glass fiber driven by a piezoelectric stimulator. In some experiments, sinusoidal force stimulation allowed assessment of the bundle’s stiffness during and after the iontophoretic application of EDTA. In other instances, a hair bundle was displacement-clamped and the force was recorded. In both instances, the restoration of a bundle’s mechanical properties provided an index of tip-link recovery.

## Acknowledgments

We thank Brian Fabella for consistent help with apparatus and computation and the members of our research groups for comments on the manuscript. M.T. is an alumna of the Frontiers in Life Science PhD program of Université Paris Diderot and thanks the Fondation Agir Pour l’Audition for a doctoral fellowship. P.M. was supported by French National Research Agency grants ANR-16-CE13-0015, ANR-11-LABX-0038, and ANR-10-IDEX-0001-02. R.G.A. was supported by Howard Hughes Medical Institute, of which A.J.H. is an Investigator.

## Supporting Information

### SI Materials and Methods

#### Preparation of the rat’s cochlea

The procedures were conducted at the Institut Curie and were approved by the Ethics Committee in accordance with the European and French National Regulation for the Protection of Vertebrate Animals used for Experimental and other Scientific Purposes (Directive 2010/63; French Decree 2013-118).

Experiments were performed on excised cochleae of Sprague-Dawley rats (*Rattus norvegicus*, Janvier Labs) of both sexes and 7-10 days of age. The dissection and isolation of the cochleae followed a published procedure (1)(2). After a rat had been euthanized and decapitated, the inner ears were extracted from the head. Each cochlear bone was carefully opened and the membranous cochlear duct uncoiled from the modiolus. After excision of the cochlear partition, the stria vascularis was removed and the tectorial membrane gently peeled away. An apical or middle turn of the organ of Corti was positioned under nylon fibers in an experimental chamber containing artificial perilymph (150 mM Na^+^, 6 mM K^+^, 1.5 mM Ca^2+^, 159 mM Cl^−^, 10 mM Hepes, 8 mM **d**-glucose, and 2 mM sodium pyruvate; pH 7.4; 315 mOsmol·kg^−1^). During the experiment, we used perfusion to change the hair bundles’ ionic environment to a variant (150 mM Na^+^, 6 mM K^+^, 3.3 mM Ca^2+^, 163 mM Cl^−^, 4 mM HEDTA, 10 mM Hepes, 8 mM **d**-glucose, and 2 mM sodium pyruvate) with a free Ca^2+^ concentration of 22 μM.

#### Preparation of the bullfrog’s sacculus

The procedures were conducted at The Rockefeller University and at the Institut Curie with the approval of the respective Institutional Animal Care and Use Committees.

Experiments were performed on hair cells from adult bullfrogs (*Rana catesbeiana*) of both sexes. After an animal had been euthanized, the sacculi were carefully removed by a standard protocol (3). Each saccular macula was sealed with tissue adhesive (Vetbond, 3M) across a 1 mm hole centered on a 10 mm square of aluminum foil. The foil was situated in a two-compartment chamber with the macular side of the sacculus facing upward. The lower compartment was filled with oxygenated artificial perilymph (114 mM Na^+^, 2 mM K^+^, 2 mM Ca^2+^, 118 mM Cl^−^, 5 mM Hepes, and 3 mM **d**-glucose; pH 7.4; 230 mOsmol·kg^−1^). The apical surface of the hair cells was exposed for 35 min at room temperature to 67 mg·L^−1^ of protease (type XXIV; Sigma) to loosen the otolithic membrane, which was carefully removed with an eyelash. The upper compartment was then filled with oxygenated artificial endolymph (2 mM Na^+^, 118 mM K^+^, 250 μM Ca^2+^, 118 mM Cl^−^, 5 mM Hepes, and 3 mM **d**-glucose; pH 7.4; 230 mOsmol·kg^−1^).

#### Measurement of hair-bundle position

Experiments on both preparations were conducted with similar apparatus. Each preparation was placed on an upright microscope (BX51WI, Olympus) and the hair cells were visualized with a 60X, water-immersed objective lens of numerical aperture 0.9 and differential-interference-contrast optics. Rat hair cells were observed during experiments with a charge-coupled-device camera (LCL-902K, Watec). Video observations of the bullfrog’s sacculus videos were conducted after an additional 4X magnification with a CMOS camera (DCC3240M, Thorlabs) or a high-speed video camera (ZYLA-5.5-CL10-W, Andor).

To record a hair bundle’s position, the preparation was illuminated with a 630 nm light-emitting diode (UHP-T-SR, Prizmatix) and the resultant shadow was projected onto a dual photodiode at a magnification of 1300X. The output of the photodiode was low-pass filtered at 2 kHz with an eight-pole anti-aliasing filter (Benchmaster 8.13, Kemo). The photodiode was calibrated by translating the bundle’s image through a succession of 10 μm steps with a mirror mounted on a piezoelectric actuator (PA 120/14 SG, Piezosystem Jena). Digital data samples were acquired at intervals of 200 μs.

#### Mechanical stimulation with fluid jets

Because they are complexly shaped and poorly cohesive, hair bundles from outer hair cells of the rat’s cochlea are difficult to stimulate with glass fibers. We therefore deflected each bundle with a fluid jet driven by a piezoelectric disk, which recruited all the stereocilia (1). When viewed under the objective lens of the microscope in the plane of the sensory epithelium, the tip of each pipette was positioned along the axis of mirror symmetry of each hair bundle at a 8 μm distance on the bundle’s abneural side. Liquid exiting the pipette therefore displaced the stereocilia towards their shortest row.

#### Mechanical stimulation with flexible fibers

Owing to the strong attachments among the stereocilia of a hair bundle from the bullfrog’s sacculus, force applied to the kinocilium uniformly displaces all the stereocilia (4). We accordingly used a flexible glass fiber attached to the kinociliary bulb to mechanically stimulate the hair bundle.

Each flexible fiber was displaced by a piezoelectric actuator (PA 4/12, Piezosystem Jena) positioned with a micromanipulator (MP-285, Sutter Instruments) and driven by an amplifier (ENV 800, Piezosystem Jena). The fiber was forged from a borosilicate capillary (1B120F-3, World Precision Instruments). After the capillary had been tapered with an electrode puller (P-2000, Sutter Instruments), its tip was melted with a platinum filament and pulled laterally with a 120 V solenoid to form a 90° angle to the shaft. The resultant fiber was approximately 100 μm in length and 1 μm in diameter. The fiber was sputter-coated with gold-palladium (Hummer 6.2, Anatech) to increase its optical contrast. To enhance the coupling of the stimulus fiber to the kinociliary bulb, we submerged the fiber’s tip in a droplet of 200 mg·L^−1^ concanavalin A for 15 min before an experiment.

Each fiber’s stiffness and drag coefficient were estimated by measuring the Brownian motion of its tip in water. We then obtained parameter values by fitting the power spectrum of the displacement to a Lorentzian relation (5). The fibers in this study had stiffnesses of 160-380 μN·m^−1^ and drag coefficients of 150-290 nN·s·m^−1^; they behaved as first-order, low-pass filters with cut-off frequencies near 200 Hz.

#### Displacement clamping

We used negative feedback to control the position of a hair bundle according to a computer-generated external command (1, 6, 7). By doing so, we were able to monitor the force required to hold a hair bundle stationary or to deflect it to a desired position. The computer’s sampling interval of 200 μs set an upper limit on the potential frequency response of the system, but a eight-pole, low-pass Bessel filter (Benchmaster 8.07, Kemo) imposed a cutoff at 2 kHz between the computer’s output and the stimulator’s input to ensure stability.

Use of the displacement-clamp system and sinusoidal stimulation allowed us to measure the decrease and subsequent recovery of hair-bundle stiffness with good temporal resolution. However, this approach confronted an inevitable problem: because the response time of the clamp system is finite, responses of progressively higher frequency become progressively less well clamped. The clamp’s settling time constant was generally about 2 ms, which corresponded to a corner frequency near 80 Hz. By selecting a stimulus frequency of 40-50 Hz, we accepted some non-ideality in clamping in the interest of improved frequency resolution in stiffness measurements.

The force *F*_SF_ exerted by the stimulus fiber against a hair bundle was estimated from the positions of the fiber measured at its base and at its tip (8):

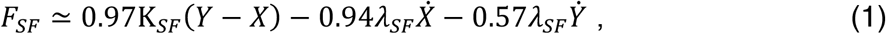

in which *K*_SF_ and *λ_SF_* represent respectively the stiffness and hydrodynamic friction coefficient of the stimulus fiber, *Y* the displacement of its base, and *X* the displacement of its tip. 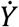 and 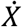 are the time derivatives of the corresponding variables. Because the stimulus frequencies were well below the cut-off frequency of the fiber, this low-frequency approximation of the periodic force applied by the fiber is expected to be accurate (8). Positive movements and forces were those directed toward a hair bundle’s tall edge.

The stiffness *K*_HB_ of each hair bundle was estimated by measuring the average force *F*_SF_ and displacement *X*_HB_ for 21 successive periods of sinusoidal stimulation. The stiffness was then computed for each sinusoidal train as

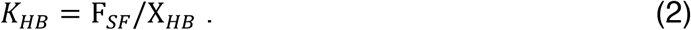

#### Voltage-clamp recording

We recorded mechanoelectrical-transduction currents of outer hair cells of the rat cochlea with whole-cell, tight-seal electrodes. Each micropipette was pulled (P^−^97, Sutter Instruments) from a thick-walled capillary (1B150F-4, WPI) and fire-polished to obtain a tip 2-3 μm in diameter. The electrode was filled with intracellular solution (142 mM Cs^+^, 11 mM Na^+^, 3.5 mM Mg^2+^, 149 mM Cl^−^,1 mM EGTA, 5 mM ATP, 0.5 mM GTP, and 10 mM Hepes; pH 7.3; 295 mOsmol·kg^−1^) and contained a chlorinated silver electrode. When immersed in standard saline, the micropipette had a resistance of 1.5-4 MΩ. The voltage across each hair cell’s membrane was controlled and currents were recorded with an amplifier (Axopatch 200B, Axon Instruments). The cell was held at a potential of −80 mV. The voltage offset was corrected before forming a gigaohm seal with a cell and the pipette’s capacitance was compensated to achieve a cut-off frequency of 1-9 kHz. Current signals were low-pass filtered at 1.25-12.5 kHz and sampled at intervals of 40-400 μs.

#### Iontophoresis

We used iontophoretic pulses to deliver Ca^2+^ chelators in the vicinity of the hair bundles. Microelectrodes were fabricated from borosilicate glass capillaries (TW 120-F, World Precision Instruments) with an electrode puller (P-80/PC, Sutter Instruments) and filled with 500 mM EDTA in 1 M NaOH. We used a current amplifier (Axoclamp 2B, Axon Instruments) to control the release of EDTA. A holding current of 10 nA kept EDTA from diffusing into the endolymphatic solution, and pulses of −100 nA released the chelator. The electrodes’ tips were directed at and situated about 2 μm from tops of the hair bundles.

## SI Notes

### Note S1. Negative hair-bundle movement during exposure to Ca^2+^ chelators

The sequence of hair-bundle forces associated with the breaking and regeneration of tip links reveals unexpected complexity in recordings from bullfrog hair cells. In six of the seven cells, there was a sustained positive offset of 20.1 ± 7.0 pN (mean ± SEM) at the end of the stimulation protocol with respect to the value before EDTA exposure (Fig. 3*A*; *SI Appendix*, Fig. S4). This force offset was absent when tip links were not broken.

In principle, this tensioning of the hair bundle would be compatible with increased activity of the adaptation machinery (7, 9). A decrease in the cytoplasmic Ca^2+^ concentration after tip-link rupture would cause the adaptation motors to ascend in the stereocilia and thus generate a negative offset in the position of the hair bundle after tip-link recovery. Nevertheless, this effect was probably masked by the presence of another, more intriguing phenomenon: a negative movement of the hair bundle that occurred seconds after tip-link breakage.

Upon exposure to Ca^2+^ chelator there was a sudden increase in the force that reflected a rise in tip-link tension, followed by the abrupt decrease that resulted from tip-link rupture. Although these observations accorded with previous studies (8, 14, 15)(10), the traces also revealed a subsequent rebound in the force (Fig. 3*A*). The force exerted by the fiber indicated that the displacement clamp acted to counter a negative movement of the hair bundle (*SI Appendix*, Fig. S4).

Although never observed in outer hair cells from the rat’s cochlea, this unexpected effect was present to a certain degree in most recordings from the two-compartment preparation of the bullfrog’s sacculus. Because this preparation recreated the environment in which hair cells normally operate, it differed from the homogeneous ionic environment of previous investigations of tip-link breakage (1, 6, 10, 11). To determine whether the unreported negative movement was consistently associated with tip-link rupture in this preparation, we measured the position of the top stereociliary row and kinocilium in unrestrained, oscillating hair bundles and applied iontophoretic pulses of Ca^2+^ chelator of various durations. Although some hair bundles responded to Ca^2+^ sequestration with canonical dynamics—a rapid negative twitch followed by a large positive displacement to a stable level—others showed a negative rebound in position (*SI Appendix*, Fig. S4). Nine of 18 spontaneously oscillating bundles displayed some degree of negative movement, ranging from −5 nm to −485 nm and averaging −166 ± 70 nm (mean ± SEM). For an additional 15 of 31 quiescent bundles that displayed negative movements, the magnitude averaged −183 ± 36 nm (mean ± SEM). This effect was most prominent when the duration of the iontophoretic pulse exceeded a few seconds, during which the negative displacement reached a plateau that variously lay either positive or negative to the bundle’s initial position. After reaching a stable plateau, the hair bundle never returned to its initial position. Moreover, because the bundle displayed a reduced stiffness and never responded to another epoch of chelator iontophoresis, the phenomenon did not result from recovery and tensioning of the tip links.

The negative hair-bundle movement often observed after tip-link disruption by Ca^2+^ chelation—or the corresponding positive force measured under displacement-clamp conditions—remains to be explained. One possibility is that the cuticular plate deforms in such a way as to alter the forces within the stereociliary cluster. For hair cells of the bullfrog’s sacculus, the cuticular plate is concave upward, a configuration that pushes the stereociliary tips together (12, 13). If the curvature of the cuticular plate were to increase after tip-link breakage, the stereocilia of the longest rank would be expected to undergo a negative displacement.

### Note S2. Lack of contribution of the kinocilium to negative movements

Aside from those associated with tip links, what other forces might act on a hair bundle? Each bundle possesses a single kinocilium that bears an axoneme with dynein motors (14). Because the kinocilium can be motile (15)(16), it might exert a force that affects the hair bundle’s position. To test that possibility, we separated the kinocilium from an oscillating hair bundle and usually held its tip several micrometers away from the stereociliary cluster with a glass microelectrode (17). Because the optical contrast of a bundle with a detached kinocilium was too low to allow the use of a photodiode, we used video microscopy to record the position of the hair bundle and a tracking algorithm (18) to trace independently the positions of the tallest stereociliary row and of the kinocilium (*SI Appendix*, Fig. S5*A*). We were then able to measure both displacements before, during, and after breaking the tip links with EDTA. Even with the kinocilium separated from and moving independently of the stereociliary cluster, four of the five hair bundles tested displayed a negative movement following EDTA exposure (*SI Appendix*, Fig. 5*B*). In some instances, the negative motion proceeded in rapid steps of irregular size, a phenomenon that occurred even when the dissociated kinocilium was immobilized against the epithelial surface by a microelectrode (*SI Appendix*, Fig. 5C). The negative hair-bundle movements thus stem from a source other than the kinocilium.

### SI Figures and Legends

**Fig. S1.**
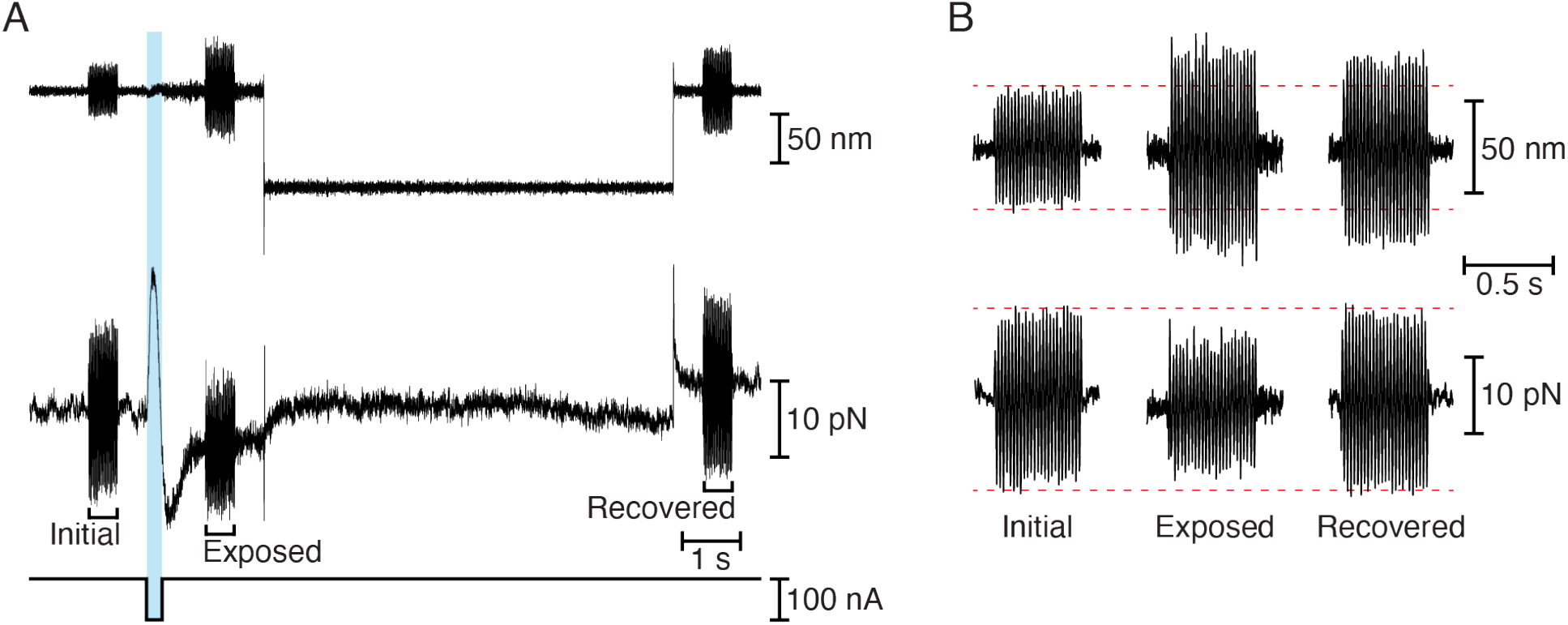
Step protocol for facilitating tip-link recovery in bullfrog saccular hair cells. (*A*) The displacement-clamp protocol imposed a step displacement of the bundle (first trace) following the iontophoretic pulse (third trace). The force necessary to clamp the bundle (second trace) diminished after iontophoresis but recovered almost completely by the experiment’s end. At three times a 500 ms epoch of ±25 nm, 50 Hz sinusoidal stimulation was superimposed on the displacement-command signal. To display the meaningful parts of the data at an appropriate scale, the transient upstrokes and downstrokes at the onset and offset of the force step have been reduced. (*B*) Enlarged records of hair-bundle displacements (top traces) and clamp forces (bottom traces) during sinusoidal stimulation highlight the phenomenon of diminished and recovered hair-bundle stiffness.

**Fig. S2.**
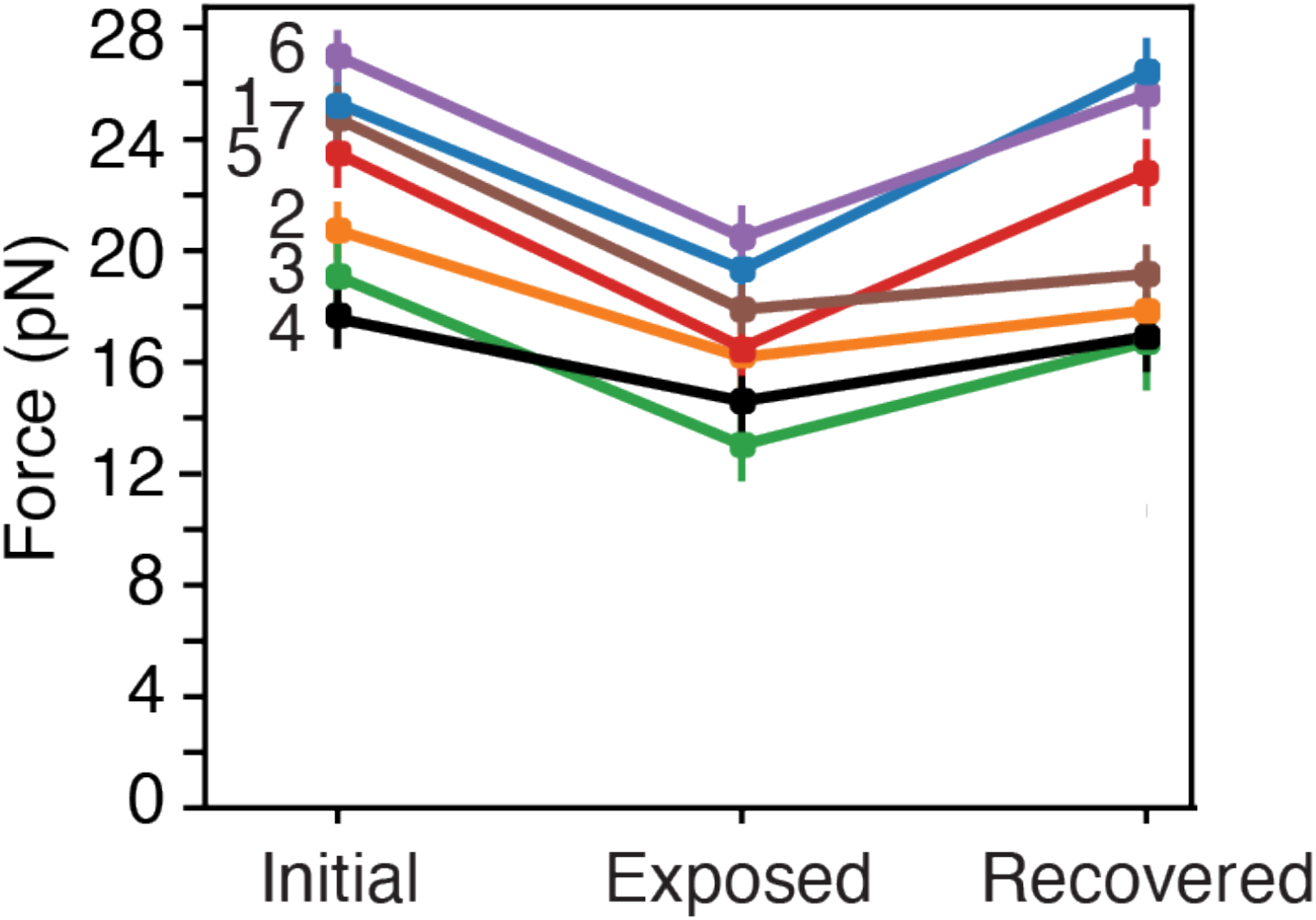
Forces measured on seven cells from the bullfrog’s sacculus during displacement-clamp measurements. In each instance, the bundle was driven sinusoidally through a distance of ±30 nm before, immediately following, and at least 6 s after the iontophoretic pulse. The force provided by the clamp is shown for bundles held in their resting positions (*Initial*), following the application of EDTA (*Exposure*), and at the experiment’s end (*Recovered*). The data show a significant decrease (*P*< 0.01 by a single-sided paired *t*-test) in the force necessary to move the bundle after chelation, followed by a significant recovery (*P* < 0.05 by the same test) towards the initial value.

**Fig. S3.**
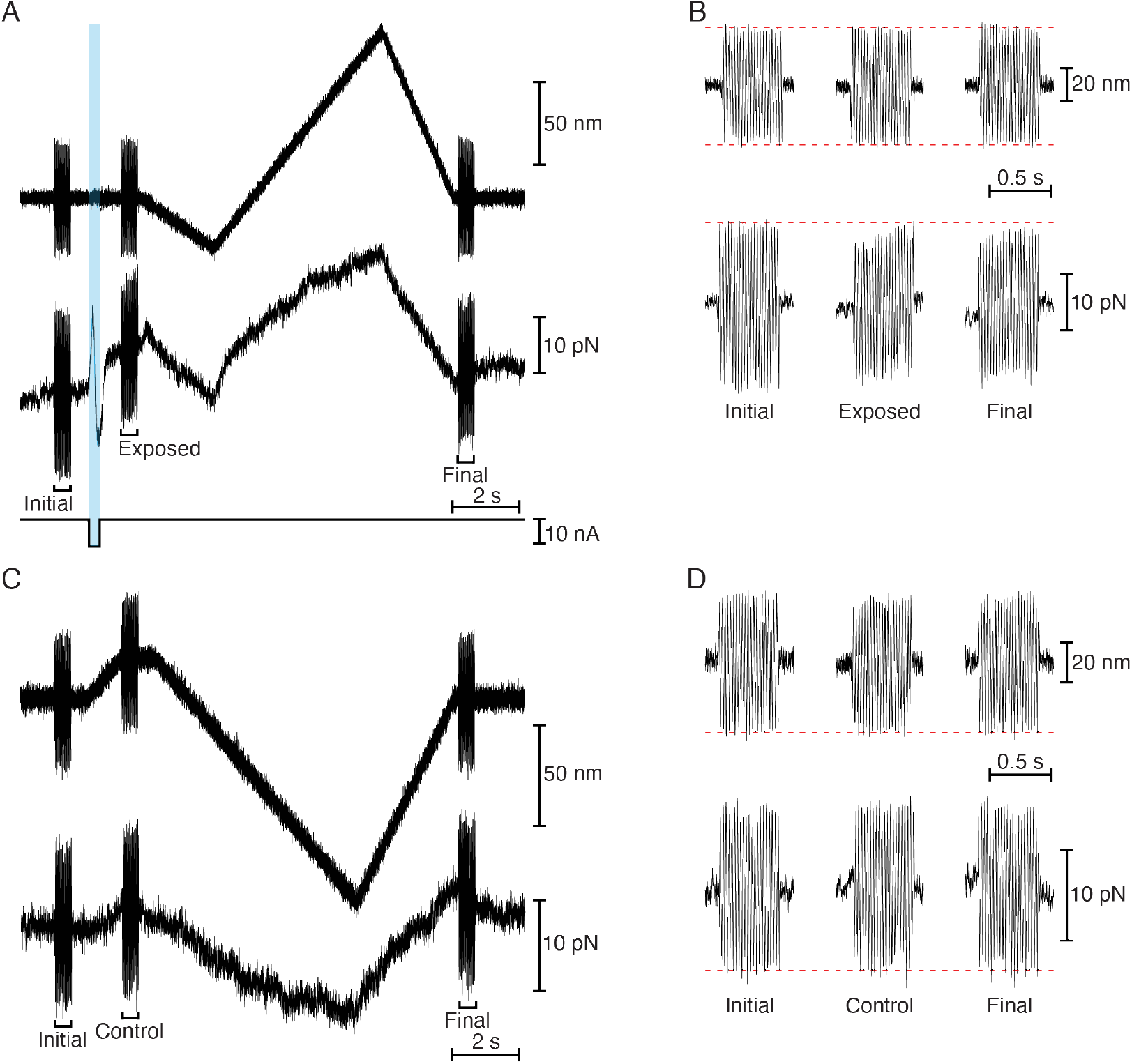
Control experiments without tip-link recovery in hair bundles of the bullfrog’s sacculus. (*A*) In a displacement-clamp protocol with a biphasic, predominantly positive displacement ramp (top trace) following the EDTA pulse (bottom trace), the force (middle trace) necessary to clamp the hair bundle to the desired position at the outset (*Initial*) decreased after iontophoresis (*Exposed*), but displayed no recovery after the ramp (*Final*). (*B*) Enlarged records of hair-bundle position (top traces) and force (bottom traces) confirm the decrease in hair-bundle stiffness and the failure of recovery after a positive ramp. (*C*) A hair bundle’s position (top trace) and force (bottom trace) during sinusoidal stimulation (*Initial*) revealed no decrease in the stiffness in the absence of iontophoresis (*Control*) or after the ramp (*Final*). (*D*) Enlarged records of hair-bundle position (top traces) and force (bottom traces) reveal no change in hair-bundle stiffness in the absence of an iontophoretic pulse.

**Fig. S4.**
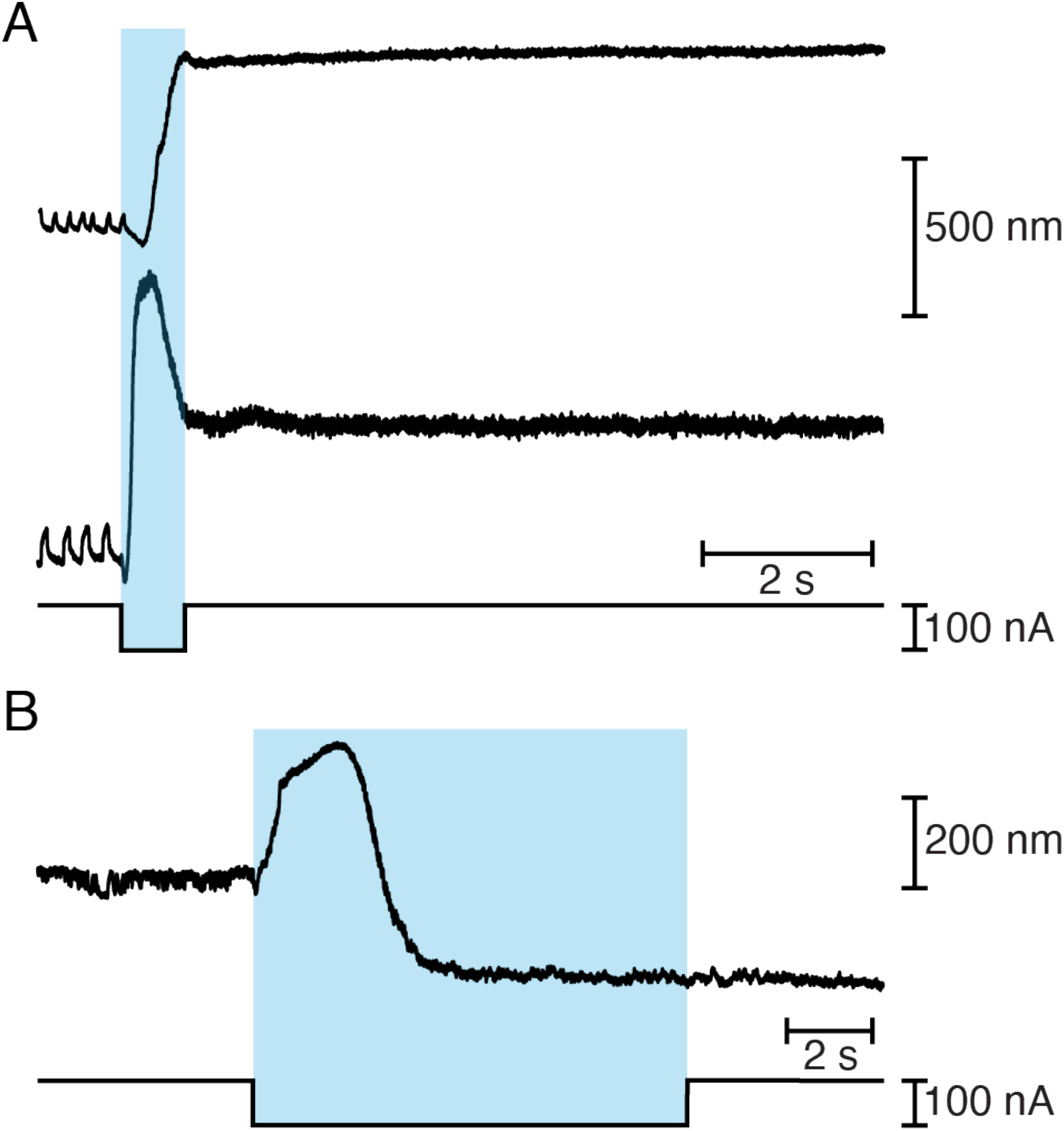
Negative hair-bundle movements after Ca^2+^ chelation in the bullfrog’s sacculus. (*A*) Two unrestrained, oscillating hair bundles displayed distinct responses to Ca^2+^ chelation. After a brief negative transient, one bundle (top trace) remained stationary at a large positive offset. The second hair bundle (bottom trace) initially followed a similar trajectory, but then underwent a sustained movement back in the negative direction. (*B*) In a similar experiment with a longer exposure to Ca^2+^ chelator, a bundle displayed a large negative movement after the initial positive movement and reached a plateau while iontophoresis was still in progress.

**Fig. S5.**
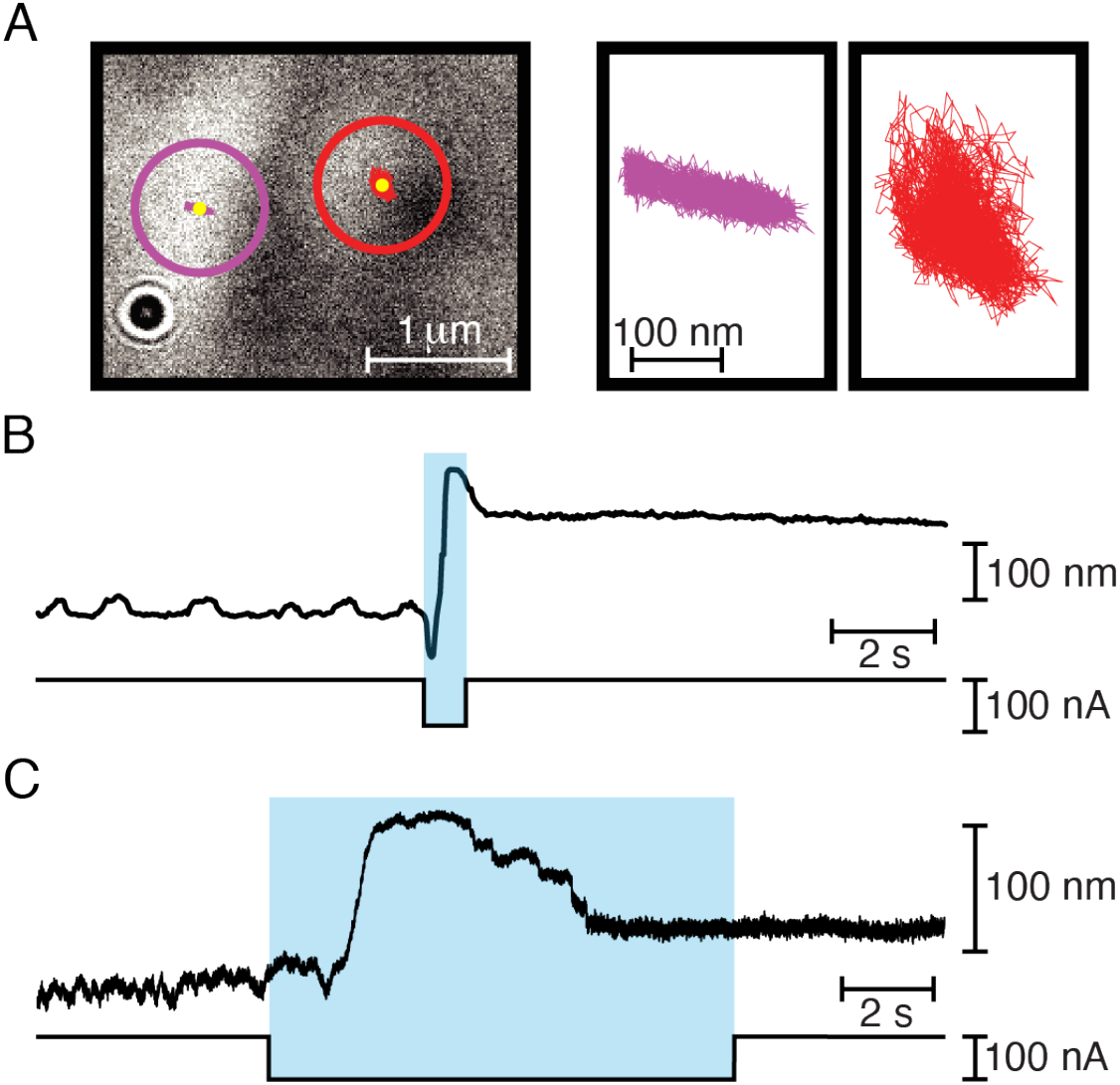
Effect of dissecting kinocilia from bullfrog saccular hair bundles. (*A*) A panel from a video record (left panel) shows the top of a hair bundle whose kinocilium had been dissected free of the stereociliary cluster. The purple circle marks the area in which stereociliary motion was tracked for 10 s at 500 frames per second and the red circle the corresponding area for the kinociliary bulb. The trajectories of the respective centroids are shown under the yellow dots at the centers of the circles. Enlarged trajectories (right panels) demonstrate that the stereocilia (purple) continued to oscillate along the bundle’s axis of mirror symmetry, whereas the kinocilium (red) underwent random motion. The scale bar at the right applies to both panels. (*B*) A record of 19 s of tracking at 30 frames per second (top trace) reveals the trajectories of a stereociliary cluster after kinociliary dissection. The rupture of tip links by iontophoresis of EDTA (bottom trace) elicited a conventional bipartite response followed by a negative movement. (*C*) In a similar experiment, the kinocilium was not only separated from the stereociliary cluster, but also held against the epithelial surface with a microelectrode. In this instance the negative motion occurred in several discrete steps, a phenomenon observed only after kinociliary dissection.

**Fig. S6.**
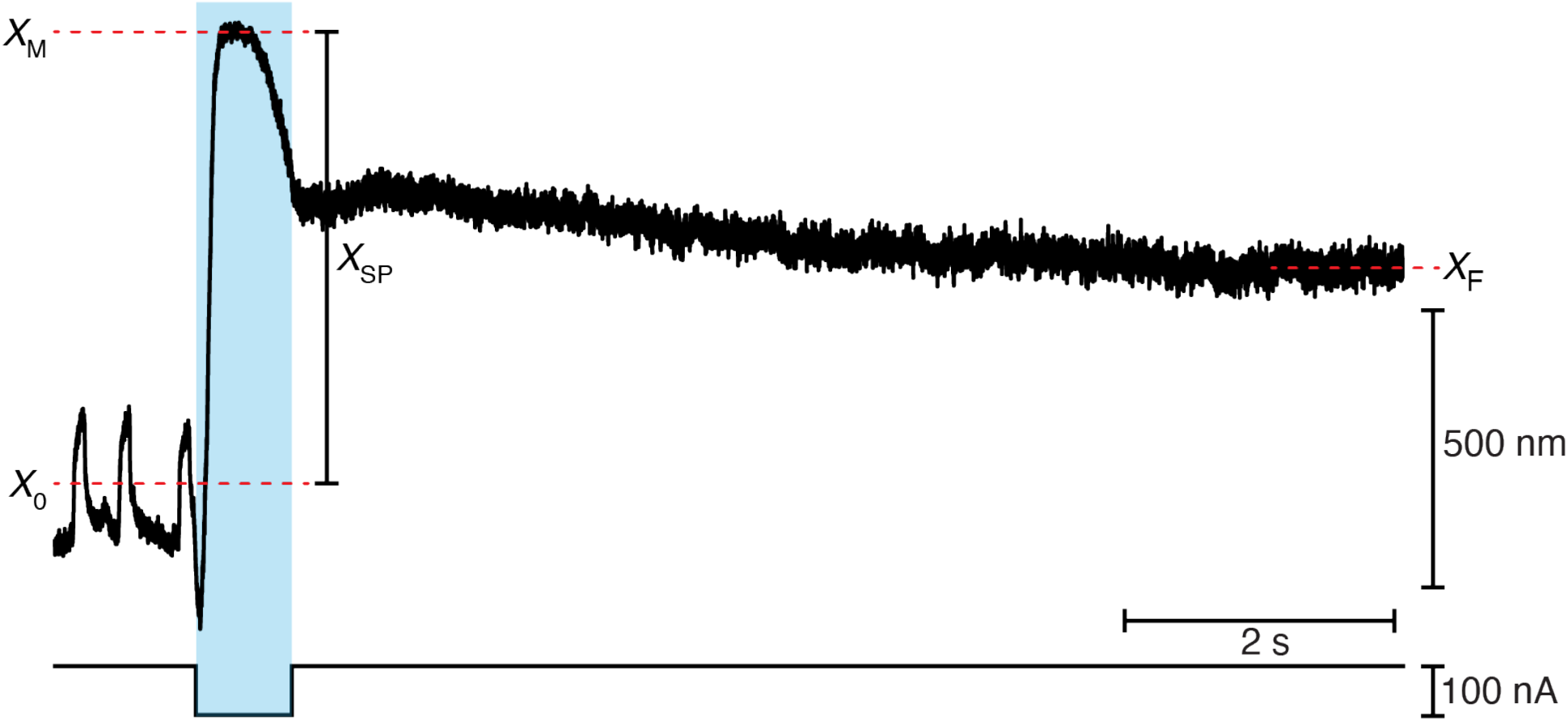
Estimation of resting tip-link tension and negative movement in hair bundles from the bullfrog’s sacculus. In a representative trace of an unrestrained, oscillating hair bundle, the bundle’s movement upon exposure to Ca^2+^ chelator (*X*_SP_) was measured from the midpoint (*X*_0_) between the maxima and minima of the spontaneous oscillations to the maximal excursion (*X*_M_) during the iontophoretic step. The final position (*X*_F_) represented the average position of the hair bundle over the last second of the experiment.

### SI Video Caption

**Video S1**. Recovery of oscillations after iontophoresis of a Ca^2+^ chelator. Viewed from above, a hair bundle from the bullfrog’s sacculus displays low-frequency spontaneous oscillations. When EDTA is expelled from the pipette at the upper left, the bundle jumps in the positive direction, to the right, and ceases to move. After the metal-coated stimulus fiber at the upper right applies force in the negative direction and is then withdrawn, the bundle resumes oscillations indicative of an intact transduction process.

## References

1. A. J. Hudspeth, Integrating the active process of hair cells with cochlear function. Nat. Rev. Neurosci. 15, 600–614 (2014).

2. T. Reichenbach, A. J. Hudspeth, The physics of hearing: fluid mechanics and the active process of the inner ear. Rep. Prog. Phys. Phys. Soc. G. B. 77, 076601 (2014).

3. D. Ó Maoiléidigh, A. J. Ricci, A Bundle of Mechanisms: Inner-Ear Hair-Cell Mechanotransduction. Trends Neurosci. 42, 221–236 (2019).

4. J. Howard, A. J. Hudspeth, Mechanoelectrical transduction by hair cells. Ann Rev Biophys Biophys Chem 17, 99–124 (1988).

5. R. A. Jacobs, A. J. Hudspeth, Ultrastructural correlates of mechanoelectrical transduction in hair cells of the bullfrog’s internal ear. Cold Spring Harb. Symp. Quant. Biol. 55, 547–61 (1990).

6. A. S. Kozlov, T. Risler, A. J. Hudspeth, Coherent motion of stereocilia assures the concerted gating of hair-cell transduction channels. Nat. Neurosci. 10, 87–92 (2007).

7. K. D. Karavitaki, D. P. Corey, Sliding adhesion confers coherent motion to hair cell stereocilia and parallel gating to transduction channels. J. Neurosci. 30, 9051–9063 (2010).

8. M. Tobin, A. Chaiyasitdhi, V. Michel, N. Michalski, P. Martin, Stiffness and tension gradients of the hair cell’s tip-link complex in the mammalian cochlea. eLife 8, e43473 (2019).

9. R. A. Eatock, D. P. Corey, A. J. Hudspeth, Adaptation of mechanoelectrical transduction in hair cells of the bullfrog’s sacculus. J. Neurosci. 7, 2821–2836 (1987).

10. A. J. Hudspeth, P. G. Gillespie, Pulling springs to tune transduction: adaptation by hair cells. Neuron 12, 1–9 (1994).

11. D. Choudhary, et al., Structural determinants of protocadherin-15 elasticity and function in inner-ear mechanotransduction. bioRxiv, 695502 (2019).

12. B. Kachar, M. Parakkal, M. Kurc, Y. D. Zhao, P. G. Gillespie, High-resolution structure of hair-cell tip links. Proc. Natl. Acad. Sci. U. S. A. 97, 13336–13341 (2000).

13. T. F. Bartsch, et al., Elasticity of individual protocadherin 15 molecules implicates tip links as the gating springs for hearing. Proc. Natl. Acad. Sci. U. S. A. 116, 11048–11056 (2019).

14. P. Kazmierczak, et al., Cadherin 23 and protocadherin 15 interact to form tip-link filaments in sensory hair cells. Nature 449, 87–91 (2007).

15. M. Sotomayor, K. Schulten, The allosteric role of the Ca2+ switch in adhesion and elasticity of C-cadherin. Biophys. J. 94, 4621–4633 (2008).

16. M. Sotomayor, W. A. Weihofen, R. Gaudet, D. P. Corey, Structure of a force-conveying cadherin bond essential for inner-ear mechanotransduction. Nature 492, 128–132 (2012).

17. J. A. Assad, G. M. G. Shepherd, D. P. Corey, Tip-link integrity and mechanical transduction in vertebrate hair cells. Neuron 7, 985–994 (1991).

18. R. Marquis, A. Hudspeth, Effects of extracellular Ca2+ concentration on hair-bundle stiffness and gating-spring integrity in hair cells. Proc. Natl. Acad. Sci. 94, 11923–11928 (1997).

19. F. Jaramillo, A. J. Hudspeth, Displacement-clamp measurement of the forces exerted by gating springs in the hair bundle. Proc. Natl. Acad. Sci. 90, 1330–1334 (1993).

20. A. A. Indzhykulian, et al., Molecular Remodeling of Tip Links Underlies Mechanosensory Regeneration in Auditory Hair Cells. PLoS Biol. 11, e1001583 (2013).

21. P. Martin, D. Bozovic, Y. Choe, A. J. Hudspeth, Spontaneous oscillation by hair bundles of the bullfrog’s sacculus. J. Neurosci. 23, 4533–4548 (2003).

22. P. Martin, A. J. Hudspeth, Compressive nonlinearity in the hair bundle’s active response to mechanical stimulation. Proc. Natl. Acad. Sci. 98, 14386–14391 (2001).

23. J. Howard, J. F. Ashmore, Stiffness of sensory hair bundles in the sacculus of the frog. Hear. Res. 23, 93–104 (1986).

24. P. Martin, A. D. Mehta, A. J. Hudspeth, Negative hair-bundle stiffness betrays a mechanism for mechanical amplification by the hair cell. Proc. Natl. Acad. Sci. U. S. A. 97, 12026–12031 (2000).

25. E. L. Wagner, J. B. Shin, Mechanisms of Hair Cell Damage and Repair. Trends Neurosci. 42, 414–424 (2019).

26. J. O. Pickles, M. P. Osborne, S. D. Comis, Vulnerability of tip links between stereocilia to acoustic trauma in the guinea pig. Hear. Res. 25, 173–183 (1987).

27. J. M. Husbands, S. A. Steinberg, R. Kurian, J. C. Saunders, Tip-link integrity on chick tall hair cell stereocilia following intense sound exposure. Hear. Res. 135, 135–145 (1999).

28. R. Kurian, N. L. Krupp, J. C. Saunders, Tip link loss and recovery on chick short hair cells following intense exposure to sound. Hear. Res. 181, 40–50 (2003).

29. Y. D. Zhao, E. N. N. Yamoah, P. G. G. Gillespie, Regeneration of broken tip links and restoration of mechanical transduction in hair cells. Proc. Natl. Acad. Sci. 93, 15469–15474 (1996).

30. A. Lelli, P. Kazmierczak, Y. Kawashima, U. Müller, J. R. Holt, Cellular/Molecular Development and Regeneration of Sensory Transduction in Auditory Hair Cells Requires Functional Interaction Between Cadherin-23 and Protocadherin-15 (2010) https:/doi.org/10.1523/JNEUROSCI.1949-10.2010.

31. E. M. Mulhall, et al., “The Dynamic Strength of the Hair-Cell Tip Link Reveals Mechanisms of Hearing and Deafness” (Biophysics, 2019) https:/doi.org/10.1101/763847 (September 21, 2020).

32. J. A. Assad, D. P. Corey, An active motor model for adaptation by vertebrate hair cells. J. Neurosci. Off. J. Soc. Neurosci. 12, 3291–3309 (1992).

## References

1. M. Tobin, A. Chaiyasitdhi, V. Michel, N. Michalski, P. Martin, Stiffness and tension gradients of the hair cell’s tip-link complex in the mammalian cochlea. eLife 8, e43473 (2019).

2. H. J. Kennedy, M. G. Evans, A. C. Crawford, R. Fettiplace, Fast adaptation of mechanoelectrical transducer channels in mammalian cochlear hair cells. Nat. Neurosci. 6, 832–836 (2003).

3. J. B. Azimzadeh, J. D. Salvi, Physiological preparation of hair cells from the sacculus of the American bullfrog. J. Vis. Exp. 55380 (2017).

4. A. S. Kozlov, T. Risler, A. J. Hudspeth, Coherent motion of stereocilia assures the concerted gating of hair-cell transduction channels. Nat. Neurosci. 10, 87–92 (2007).

5. J. D. Salvi, D. Ó Maoiléidigh, B. A. Fabella, M. Tobin, A. J. Hudspeth, Control of a hair bundle’s mechanosensory function by its mechanical load. Proc. Natl. Acad. Sci. U. S. A. 112, E1000–1009 (2015).

6. F. Jaramillo, A. J. Hudspeth, Displacement-clamp measurement of the forces exerted by gating springs in the hair bundle. Proc. Natl. Acad. Sci. U. S. A. 90, 1330–1334 (1993).

7. P. Martin, A. D. Mehta, A. J. Hudspeth, Negative hair-bundle stiffness betrays a mechanism for mechanical amplification by the hair cell. Proc. Natl. Acad. Sci. U. S. A. 97, 12026–12031 (2000).

8. V. Bormuth, J. Barral, J.-F. Joanny, F. Jülicher, P. Martin, Transduction channels’ gating can control friction on vibrating hair-cell bundles in the ear. Proc. Natl. Acad. Sci. U. S. A. 111, 7185–7190 (2014).

10. J. A. Assad, G. M. G. Shepherd, D. P. Corey, Tip-link integrity and mechanical transduction in vertebrate hair cells. Neuron 7, 985–994 (1991).

11. R. Marquis, A. Hudspeth, Effects of extracellular Ca^2+^ concentration on hair-bundle stiffness and gating-spring integrity in hair cells. Proc. Natl. Acad. Sci. 94, 11923–11928 (1997).

12. A. J. Hudspeth, Mechanoelectrical transduction by hair cells in the acousticolateralis sensory system. Annu. Rev. Neurosci. 6, 187–215 (1983).

13. R. A. Jacobs, A. J. Hudspeth, Ultrastructural correlates of mechanoelectrical transduction in hair cells of the bullfrog’s internal ear. Cold Spring Harb. Symp. Quant. Biol. 55, 547–61 (1990).

14. C. Spoon, W. Grant, Biomechanics of hair cell kinocilia: Experimental measurement of kinocilium shaft stiffness and base rotational stiffness with Euler-Bernoulli and Timoshenko beam analysis. J. Exp. Biol. 214, 862–870 (2011).

15. R. E. Bowen, Movement of the so-called hairs in the ampullar organs of fish ears. Proc. Natl. Acad. Sci. U. S. A. 17, 192–194 (1931).

16. A. Rüsch, U. Thurm, Spontaneous and electrically induced movements of ampullary kinocilia and stereovilli. Hear. Res. 48, 247–263 (1990).

17. A. J. Hudspeth, R. Jacobs, Stereocilia mediate transduction in vertebrate hair cells. Proc. Natl. Acad. Sci. U. S. A. 76, 1506–1509 (1979).

18. J. Y. Tinevez et al., TrackMate: An open and extensible platform for single-particle tracking. Methods 115, 80–90 (2017).

